# Evolutionary history of phycoerythrin pigmentation in the water bloom-forming cyanobacterium *Microcystis aeruginosa*

**DOI:** 10.1101/485508

**Authors:** Yuuhiko Tanabe, Haruyo Yamaguchi

## Abstract

*Microcystis aeruginosa* is a bloom-forming cyanobacterium found in eutrophic fresh-and brackish water bodies worldwide. As typical for cyanobacteria, most *M. aeruginosa* strains are blue-green in color owing to the concomitance of two photosynthetic pigments, phycocyanin (PC) and chlorophyll *a*. Although less common, *M. aeruginosa* strains that are brownish in color owing to the presence of another pigment phycoerythrin (PE) have been documented. However, the genomic basis, phylogeny, and evolutionary origin of PE pigmentation in *M. aeruginosa* have only been poorly characterized until date. In the present study, we sequenced and characterized the genomes of five PE-containing *M. aeruginosa* strains. Putative PE synthesis and regulation genes (the *cpe* cluster) were identified in all five sequenced genomes as well as in three previously published *M. aeruginosa* genomes. Of note, Absorption spectra indicated that the PE content, but not PC content, was markedly altered in response to availability of red/green light in all PE-containing strains. This was consistent with the presence of *ccaS*/*ccaR*, a hallmark of type II chromatic adapter, in the *cpe* cluster. Phylogenetic analyses of core genome genes indicated that PE-containing genotypes were located in three different phylogenetic groups. In contrast, the genomic organization of the *cpe* cluster was mostly conserved regardless of genomic background. Additionally, the phylogenies of PE genes were found to be congruent, consistent with the core genome phylogeny. A comparison of core genome and PE genes showed a similar level of genetic divergence between two PE-containing groups. These results suggest that genes responsible for PE pigmentation were introduced into *M. aeruginosa* early during evolution and were repeatedly lost thereafter possibly due to ecological adaptation. Additional horizontal gene transfer (HGT) later during evolution also contributed to the present phylogenetic distribution of PE in *M. aeruginosa*.

## Introduction

*Microcystis aeruginosa* is one of the most common water bloom-forming cyanobacteria found in fresh and brackish water bodies worldwide (Harke et al., 2016). Bloom occurrence of *M. aeruginosa* in lakes and reservoirs causes several environmental problems including foul odor and bottom-layer hypoxia. However, the production of hepatotoxic cyanotoxins called microcystins can be regarded as a problem of greatest concern (Harke et al., 2016). Cases of intoxication of human (Jochimsen et al., 1998), livestock (Beasley et al., 1989) and wild animals (Miller et al., 2010a) by the microcystin contaminations have been sporadically reported around the globe.

In almost all cyanobacteria including *M. aeruginosa*, a water-soluble macromolecular complex, the phycobilisome is present associated with thylakoid membranes, and serves as an apparatus that complements the light-harvest function of photosystem II (PSII). The key function of phycobilisomes enables the cells to absorb light within the 550–600 nm wavelength range which chlorophyll molecule cannot absorb and thereby it facilitates the photosynthetic growth of cells where light of longer wavelength is not available (e.g., deep underwater environments) (Kirk, 1994). Phycobilisomes consist of core (allophycocyanin, APC) and rod phycobiliproteins such as phycocyanin (PC) and phycoerythrin (PE), and linker polypeptides that unite the cores and rods (Watanabe and Ikeuchi, 2013). The type and the number of rod phycobiliproteins incorporated into the phycobilisome mainly determines the color of the cyanobacterial cells (Six et al., 2007). Several cyanobacterial species or strains are known to be capable of changing the color in response to availability of different wavelengths of light, by, for example, replacing the rod phycobiliprotein with other phycobiliproteins. This process is called chromatic adaptation (CA) (Tandeau de Marsac, 1977; Montgomery, 2017). CA can be categorized on the basis of the different modes of phycobiliprotein accumulation. Type I (CA1) strains show no change in the content of both PE and PC; Type II (CA2) strains show an increase in PE but not in PC content under green light; type III (CA3) strains increase both PC and PE content under red and green light, respectively (Tandeau de Marsac, 1977). Other complex types of CA include a change in phycobilin content in the chromophore in response to blue/green light (CA4), and dynamic remodeling of phycobilisomes under far-red light perception (FaRLiP) (Montgomery, 2017).

Like other cyanobacteria, most *M. aeruginosa* strains are characterized by a blue-green color owing to the presence of photosynthetic pigments: chlorophyll *a* and PC (Otsuka et al., 1998a). *M. aeruginosa* strains with grayish-to dark brown colors have also been isolated, albeit rarely (Otsuka et al., 1998a; Schatz et al., 2007; Jeong et al., 2018). Previous studies have suggested that the brownish color of *M. aeruginosa* can be attributed to the presence of another phycobiliprotein PE in addition to PC and chlorophyll *a* (Otsuka et al. 1998a; Schatz et al., 2007). Previous 16S rDNA and ITS phylogenetic analyses have indicated that PE-containing strains are genetically highly similar to PE-deficient strains of *M. aeruginosa* (Otsuka et al., 1998b), representing an intraspecific diversity rather than difference in species (Otsuka et al., 1999). However, the PE pigmentation of *M. aeruginosa* has thereafter never been characterized in a phylogenetic framework. This may be partly because of the unavailability of PE-containing strains sufficient for in-depth phylogenetic analyses and inferences. Additionally, genes for PE synthesis and regulation in *M. aeruginosa* have not been described in detail. It is also unknown whether PE-containing *M. aeruginosa* strains regulate their PE content in response to different light wavelengths and undergo CA.

To address these questions, we characterized five brownish colored *M. aeruginosa* strains from Japanese lakes using whole genome sequencing, phylogenetic, and pigment analyses. Our data revealed that PE-containing strains were not monophyletic. PE synthesis, regulation and genomic architecture was found to be highly conserved among PE-containing *M. aeruginosa* strains. We have discussed the possible evolutionary history of PE pigmentation in *M. aeruginosa* within the scope of this study.

## Materials and methods

### Strains

*Microcystis aeruginosa* Ks07TS04 was isolated from a bloom sample collected in Lake Kasumigaura, Japan (N36° 4’ 46”, E140° 12’ 32’’) on August 3, 2007, using a micropipetting method. Additionally, the strains Ki05YA01, Tn05Ak01, (Tanabe et al., 2009), Ks07TS139, Hs07SP05 (Tanabe et al., 2011), and NIES-1211 (Tanabe et al., 2018) were used for subsequent analyses. All six strains appeared brownish in culture under white light, indicative of the presence PE (Figure 1, Supplementary Figure S1). The former five strains were deposited at the Microbial Culture Collection of the National Institute for Environmental Studies (MCC-NIES, Tsukuba, Japan) under strain no. NIES-4264, NIES-2519, NIES-2520, NIES-2521, and NIES-2522, respectively. Additionally, a PE-deficient strain *M. aeruginosa* NIES-4234 (=Sj) (Tanabe et al., 2018) was used as a reference for spectral analysis of phycobiliproteins. Other strains used in PCR detection of PE are listed in Supplementary Table S1.

**Fig. 1.**
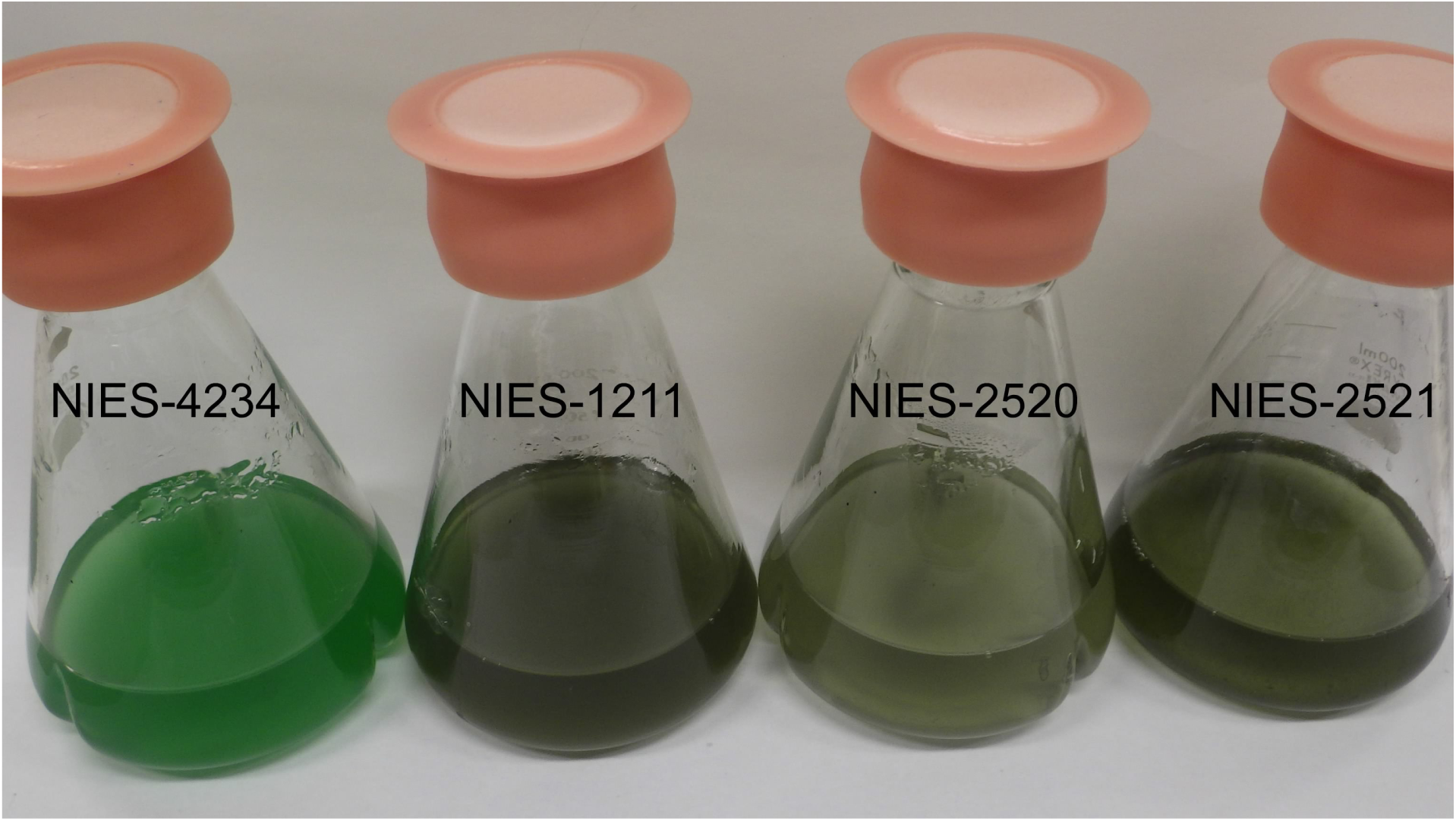
Culture of *M. aeruginosa* strains under white light. The phycoerythrin (PE)-containing strains NIES-1211, NIES-2520, and NIES-2521 are brownish in color, whereas the PE-deficient strain NIES-4234 is blue-green in color.

### Spectral analyses

For this purpose, 2 ml of mid-exponential cultures of *M. aeruginosa* grown in MA medium under white light (Kasai et al., 2004) was inoculated into 2 ml of fresh MA medium. Samples were grown in 20 ml screw cap test tubes at 23–25 °C under white, green, and red lights (24 hr, 10 μmol photons m^−2^ s^−1^) for 1 week. *In vivo* absorption spectra were measured using a spectrophotometer UV-1800 (Shimadzu, Kyoto, Japan). For *in vitro* absorption spectra, 1 ml culture was sonicated for 5 min, and centrifuged at 20 000 *g* for 5 min. The pellet was suspended in 1 ml of 1x PBS buffer (pH 7.2) in a microtube. Zirconium beads (0.1 mm in diameter) were added into the cell suspension and cells were disrupted using a mini Bead-Beater (Biospec Products, Bartlesville, OK) at 4 200 rpm for 60 s, and then centrifuged at 20 000 *g* for 10 min. Supernatants were used for measurement of absorption spectra.

### Whole genome shotgun analyses

*Microcystis aeruginosa* NIES-2519, NIES-2522, and NIES-4264 were not axenic. Bacterial contaminants in these samples were removed according to a previously published protocol (Tanabe et al., 2018). Genomic DNA of each strain was extracted using NucleoBond® AXG Columns with Buffer set III (Macherey-Nagel, Düren, Germany). DNA was fragmented to approximately 450 bps using Covaris M220 (Covaris, Woburn, MA). A 450-bp fragmented library was constructed using the NEBNext Ultra DNA Library Prep Kit for Illumina (New England Biolabs, Ipswich, MA). DNA was sequenced using the MiSeq platform (Illumina, San Diego, CA) with the 500-cycle MiSeq Reagent Kit v2 and Nano Kit v2. *De novo* assembly of the contigs was performed using Spades ver. 3.10.1 (Bankevich et al., 2012) and automated annotations were performed using Prokka ver. 1.12 (Seeman, 2014). Contigs of possible contaminants were excluded on the basis of lower coverage (identified by Spades) and a BLAST analysis. The whole genome shotgun datasets of NIES-2519, NIES-2520, NIES-2521, NIES-2522 and NIES-4264 have been deposited in the DNA Database of Japan (DDBJ) under accession nos. BHVO01000000, BHVP01000000, BHVQ01000000, BHVR01000000, and BHVS01000000, respectively. The whole genome data for NIES-1211 was already published (Tanabe et al., 2018).

### Phylogenetic analyses

Multilocus sequence typing (MLST) of *M. aeruginosa* NIES-4264 was performed according to a previously published protocol (Tanabe et al., 2007, 2009). MLST data of other phycoerythrin-containing strains used in this study have been previously published (Tanabe et al., 2018), with the exception of MLST data for the phycoerythrin-containing strain *M. aeruginosa* MC19 (Jeong et al., 2018), which were retrieved from GenBank. Phylogenetic analyses of concatenated sequences of the seven MLST loci of 251 strains (Supplementary Table S1) were performed by neighbor-joining (NJ) and maximum-likelihood (ML) methods using MEGA version 6 (Tamura et al., 2013) and RAxML (Stamatakis, 2006), respectively. The NJ tree reconstruction and bootstrap analysis (1 000 resamplings) employed the maximum composite likelihood substitution model with uniform nucleotide substitution rates among sites and lineages. RAxML was run at Cipress Science Gateway (Miller et al., 2010b) under default settings, except that bootstrap analysis was performed with 1 000 replicates. The MLST alignment data for phylogenetic analyses was provided as Supplementary Data S1. Phylogenetic analyses of DNA sequences of PE genes were performed using the NJ method as described above. Amino acid alignments of CpeA and CpeB were generated using MUSCLE (Edgar, 2004) implemented in MEGA. The alignments consist of 227 and 381 amino acids for CpeA, and CpeB, respectively. Methods for phylogenetic reconstruction are identical as those for MLST, with the exception that the JTT protein substitution model, with the option of “pairwise deletion” of gaps, was used for reconstruction of NJ tree, and “protein GAMMA” was used for RAxML analyses. Gaps were removed from the alignment prior to RAxML analyses.

### Phycoerythrin (PE) gene detection

Genomic DNAs used in the previous study (Tanabe et al., 2018) were used as templates. PCR detection of one of the PE genes *cpeA* was performed using the primer pairs, cpeAF/cpeAR (Supplementary Table S2). PCR conditions and sequencing methods were the same as those reported for MLST loci (Tanabe et al., 2007) except for annealing temperatures (Supplementary Table S2).

## Results and discussion

### Phycoerythrin (PE)-containing strains of *M. aeruginosa*

We analyzed six *Microcystis aeruginosa* strains, assumed to be putative PE-containing strains based on their brownish color (Figure 1 and Supplementary Figure S1). Consistent with this visual inspection, *in vivo* absorption spectra of all, except the blue-green strain, showed a dominant peak at ≈573 nm indicative of the presence of PE (Figure 2A). The culture of the three strains that represented different phylogenetic groups (groups A, G, and K) under red light was green in color (Supplementary Figure S2). *In vitro* spectra showed a marked change in the height of the absorption peak of PE under different green/red light availability, whereas the height of the absorption peak of PC changed only slightly (Figure 2B, C, D), suggesting that all three strains represent a type II chromatic adapter (CA2, Tandeau de Marsac, 1977). These results suggest that PE-containing *M. aeruginosa* strains belonging to different lineages shared a similar genomic architecture for PE synthesis and regulation.

**Fig. 2.**
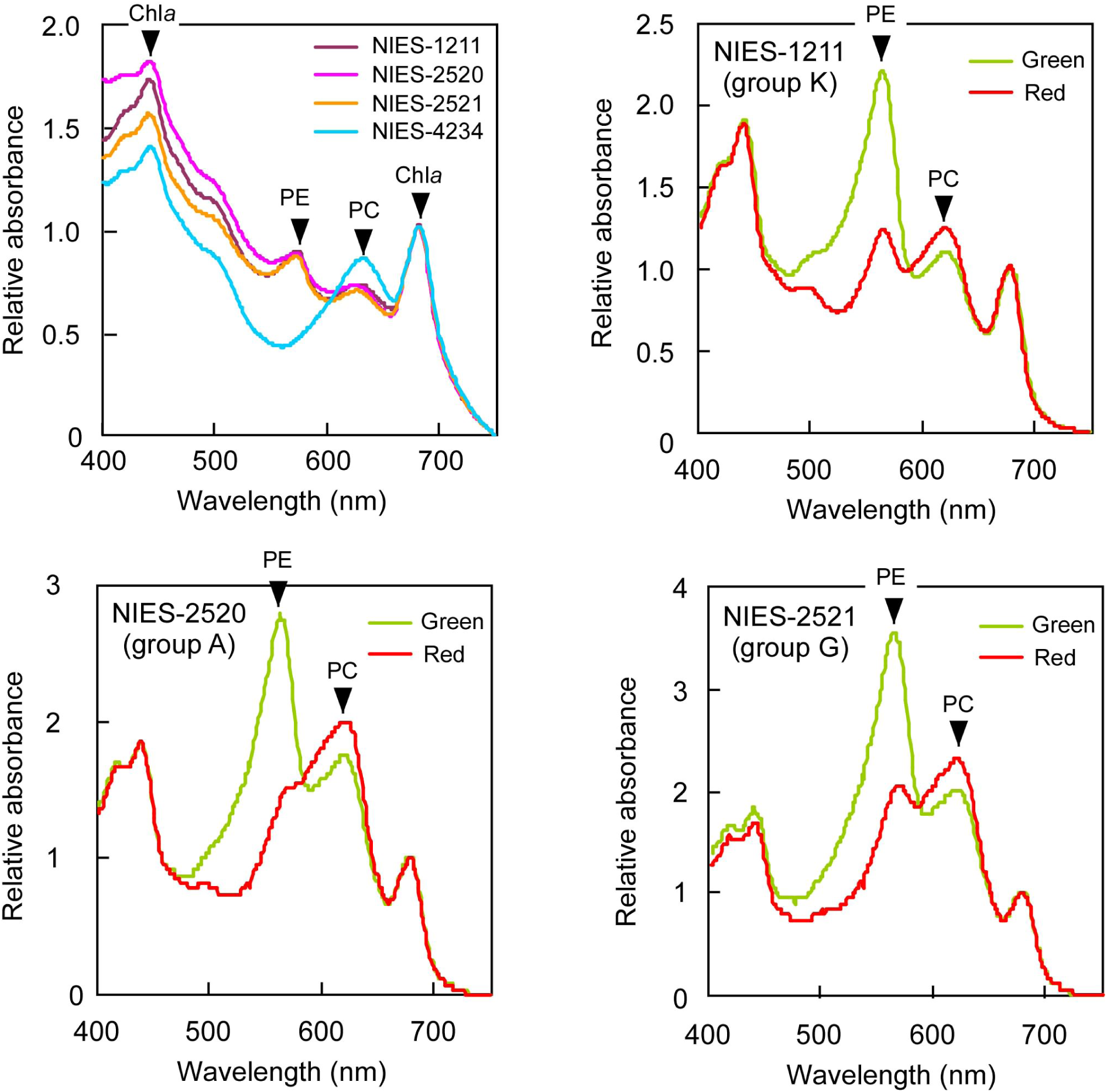
Absorption spectra. All spectra were normalized using the chlorophyll *a* absorption peak at 680 nm. **a**, *in vivo* spectra of *M. aeruginosa* strains grown under white light (full spectrum). *In vitro* spectra of NIES-1211 (**b**), NIES-2520 (**c**), and NIES-2521 (**d**), cultured under green or red light. The *in vitro* spectra also show peaks due to contamination with chlorophyll *a*.

### Phylogeny of phycoerythrin (PE)-containing strains of *M. aeruginosa*

PE-containing strains are extremely rare in *M. aeruginosa*. Among >20 *M. aeruginosa* strains for which whole genome sequence data are currently available, only three strains MC-19, NIES-1211, and T1-4 possess genes for PE synthesis, and thus possibly contain PE (Supplementary Table S1). Additionally, a comprehensive PCR survey of >200 available strains targeting one of the PE genes, *cpeA*, led to the discovery of two putative PE-containing strains, MCS3 and Ia05Yo06 (Supplementary Table S1). Phylogenetic analysis on the basis of MLST genes (representing core genome genes) indicated PE-containing strains were not monophyletic; they distributed in three different phylogenetic groups A, G and K (Figure 3). The non-monophyletic nature of PE-containing strains is consistent with an earlier ITS-based study wherein PE-containing strains were found in two different phylogenetic groups (Otsuka et al., 1999). Given the substantial intra-group genetic distance between the PE-strains in groups A and K, it is highly likely that these strains represent stable PE-containing ecotypes. On the other hand, the PE-containing NIES-2521 shares an identical *ftsZ* genotype with other PE-negative group G strains (Supplementary Table S1). This suggests the possibility that the acquisition of PE by NIES-2521 may have been a recent event.

**Fig. 3.**
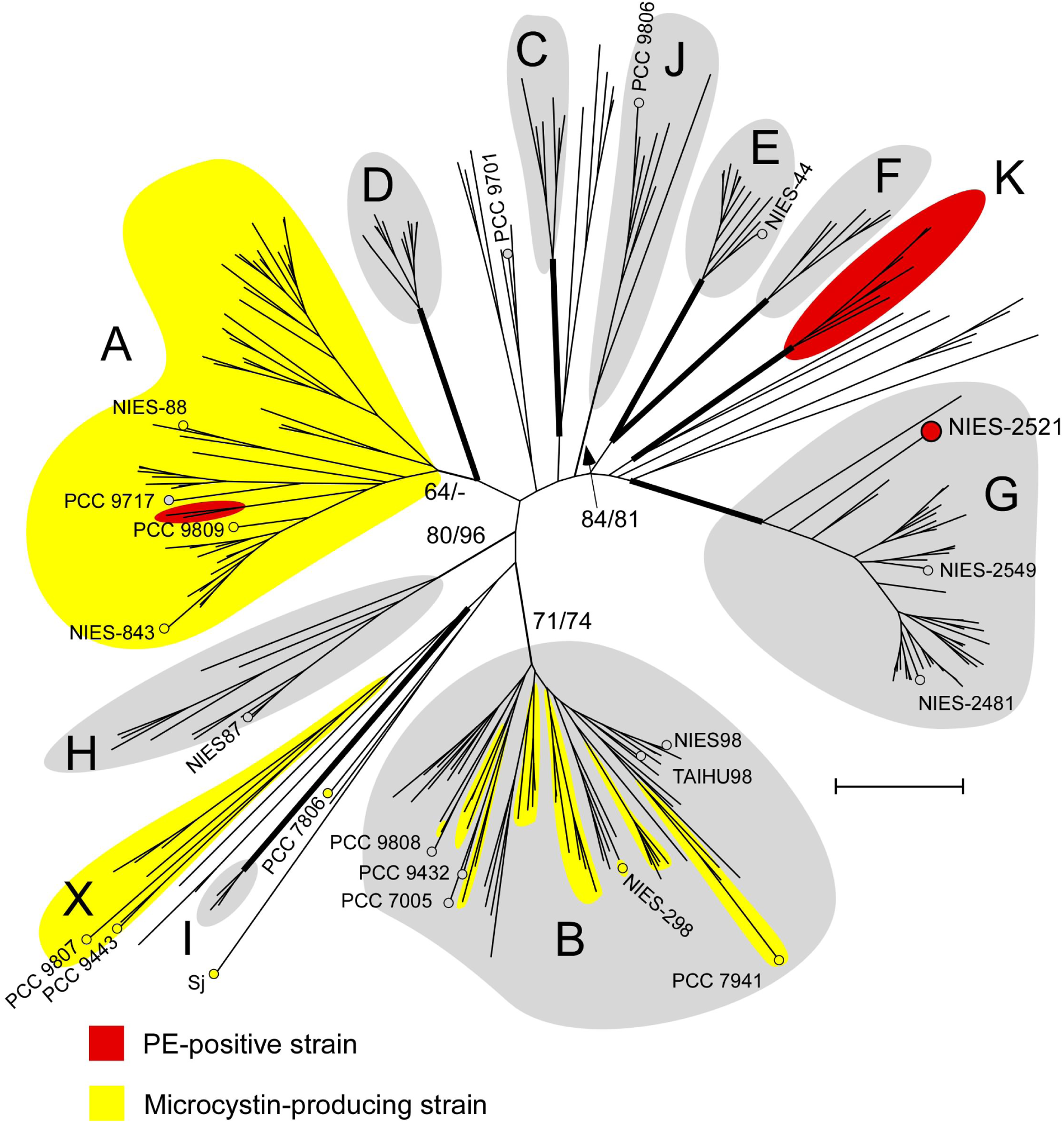
Phylogeny of PE-containing strains of *Microcystis*. A multilocus NJ phylogenetic tree was constructed on the basis of concatenation of the seven housekeeping loci (2 992 bps in total). Intraspecific phylogenetic groups were defined according to a previous report (Tanabe et al., 2018) with a newly designated group K, comprising exclusively of PE-containing strains. Numbers at the major branches indicate bootstrap values (NJ/RAxML). Thickened lines indicate branches supported by > 95% bootstrap values in both NJ and RAxML. *M. aeruginosa* strains, for which a whole genome sequence was available, are indicated by circle with the strain name (except for PE-containing strains in groups A and K, which are indicated in Figure 4 at higher magnification). Strain PCC 9717, which harbors an incomplete microcystin synthetase gene cluster (Humbert et al., 2013), is indicated in gray color. Scale bar, 0.005 substitutions per sites.

### Genomic organization of phycoerythrin (PE) genes in *M. aeruginosa*

The draft genome sequences of eight strains, including three *M. aeruginosa* strains (MC19, NIES-1211, and T1-4) whose whole genome data have been published, identified a set of putative genes for PE synthesis and regulation (hereafter designated the “*cpe* cluster”) in two to three different contigs (Figure 4, Supplementary Table S3). The available contig data indicates that the *cpe* cluster is located separately at two locations within each genome. BLAST searches targeting published *M. aeruginosa* genomes indicated that most genes in this gene cluster were exclusive to the eight PE-containing strains (Figure 4). The PE-containing strain-specific genes include genes for the alpha and beta chain of phycoerythrin (*cpeA* and *cpeB*), phycoerythrin-associated linker proteins (*cpeC* and *cpeD*), phycoerythrobilin (PEB) synthesis (*pebA* and *pebB*), phycoerythrin lyase (*cpeS, cpeT*, and *cpeZ*), and an activator for *cpe* expression (*cpeR*) (Hirose et al, 2017). Notably, a green/red light sensor histidine kinase gene (*ccaS*) and its cognate response regulator gene (*ccaR*) are shared in the cluster. Both gene products are cooperatively involved in the regulation of PE synthesis under different green/red light availability in type II chromatic adapters (CA2) (Hirose et al., 2010). This suggests that despite their different genomic background, all PE-containing *M. aeruginosa* strains share the molecular basis for CA2, which is consistent with the result observed in absorption spectra that showed a similar pattern of CA (Figure 2B, C, D). Besides *ccaS*/*ccaR*, another genes for putative sensor His-kinase (ORF3) and response regulator (ORF2) were found in the *cpe* cluster. Although the function of these proteins remains to be solved, it is clear that ORF3 is not a CcaS homologue because it lacks GAF domain that is essential for the photoreceptor function of cyanobacteriochrome (Ikeuchi and Ishizuka, 2008). The organization of genes within the *cpe* cluster in the genome of the three different phylogenetic groups is highly conserved, except for minor modifications such as the insertion/deletion of genes for putative transposases and endonucleases involved in genome rearrangement, and putative small ORFs (Figure 4). This observation suggests that the *cpe* gene clusters in *M. aeruginosa* share a same common ancestor. Unlike other cyanobacteria such as *Planktothrix* spp. (Tooming-Klunderud et al., 2013), the phycocyanin (PC) operon (*cpc* operon) was not located adjacent to the *cpe* cluster, and found scattered throughout the genome as in the case of *Geminocystis* spp. (Hirose et al., 2017).

**Fig. 4.**
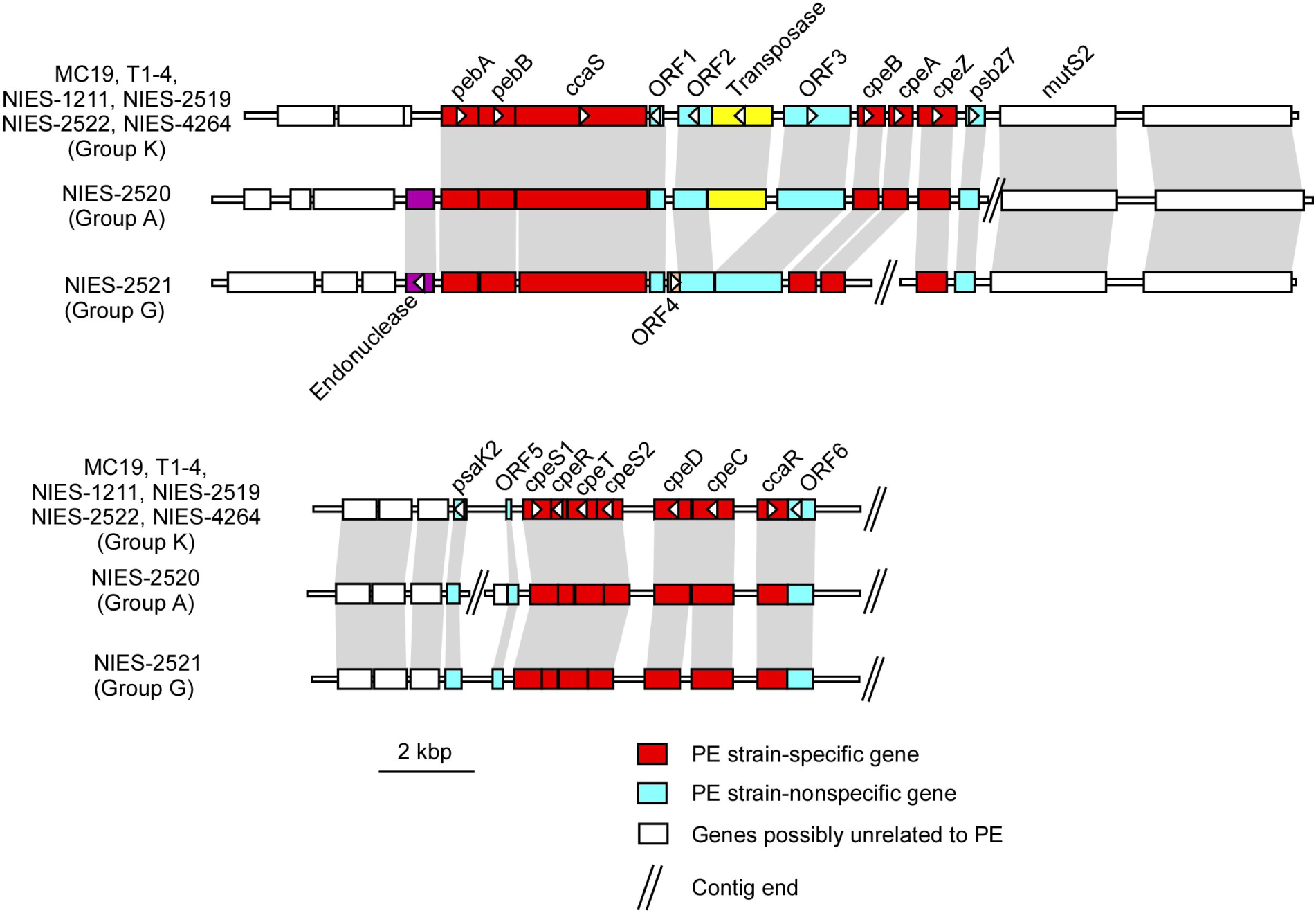
Organization of genes functioning in PE synthesis and regulation in the genome of PE-containing *M. aeruginosa* strains. Homologous genes are indicated by gray shading. ORF4, endonuclease and transposase, are not shared in all strains and are indicated in different colors. ORFs 1 to 6, whose involvement in PE regulation is not clear, are indicated in blue.

### Phylogenetic distribution and molecular evolution of PE genes

Genes necessary for adaptive traits in *M. aeruginosa* have frequently undergone *horizontal* gene transfer (HGT) (e.g., salt resistance, Guljamow et al., 2007; Tanabe et al., 2018). Owing to this, MLST phylogenetic analysis raises the possibility that HGT is responsible for the patchy distribution of PE genes in *M. aeruginosa*. In order to confirm whether this is true, phylogenetic analyses of PE genes were performed. The phylogenetic tree of a broad range of cyanobacteria (Figure 5) indicated significant discordance between the core genome (conserved proteins, Shih et al., 2013) and CpeAB phylogenies. For example, strains belonging to B2 in the core genome tree scattered across the CpeAB tree. This suggests frequent HGT of PE genes in the course of *Cyanobacteria* diversification. In the core genome tree, the strains most closely related to *M. aeruginosa* were *Cyanothece* spp. PCC 7822 and PCC 7824, both of which were distantly related to *M. aeruginosa* in the CpeAB tree. The CpeAB tree also indicated that PE genes in *M. aeruginosa* were monophyletic. These results suggest that PE genes in *M. aeruginosa* share a single origin, probably imported via HGT from another cyanobacterium. However, the identification of the donor was not possible owing to the substantial genetic divergence between *M. aeruginosa* and most closely related strains (i.e., *Leptolyngbya* sp. PCC 7376 and *Limnothrix rosea* IAM M200). All the individual PE gene phylogenies within *M. aeruginosa*, including those of regulatory genes (e.g., *ccaS*), were mostly concordant and consistent with the core genome (MLST) phylogeny (Figure 6 and Supplementary Figure S3). Two hypotheses may explain the observed pattern of PE gene divergence (Supplementary Figure S4). First, PE genes were introduced into *M. aeruginosa* via early HGT, and were repeatedly lost in the subsequent divergence into three intraspecific groups A, G, and K (“PE-early”). Second, before the intra-group divergence of groups A and K, recurrent HGT drove the spread of PE genes among *M. aeruginosa* (“PE-late”). These hypotheses were tested by comparing the nucleotide divergence between the core genome and the PE genes. The rationale behind these tests is that a smaller DNA divergence of PE genes compared with those of core genome genes must favor the “PE-late,” while similar levels of DNA divergence between the two must be consistent with the “PE-early” hypotheses. Nucleotide divergence between the core genome genes and PE genes was found to be similar when groups A and G/K were compared, while a smaller divergence of PE genes than core genome genes was observed when groups G and K were compared (Figure 7). These results suggest that the most plausible evolutionary scenario is a composite of the two hypotheses mentioned above: PE genes were imported into the common ancestor of the groups A and K via interspecific HGT, and were repeatedly lost in group A; the group G strain (NIES-2521) acquired PE genes via intraspecific HGT from group K strains (Figure 8). The PE pigment evolution in another bloom-forming cyanobacterium *Planktothrix* spp. from “green to red,” in which one PE-containing strain likely obtained the PE gene cluster via HGT, has previously been reported (Tooming-Klunderud et al., 2013). Contrastingly, our data suggests that the observed pattern of PE distribution in *M. aeruginosa* can be largely explained by the “red to green” evolution. Unfortunately, the branching order of the major groups in *M. aeruginosa* including groups A, G, and K, was unresolved due to low statistical support, and complicated an exact inference regarding import of PE genes into each intraspecific lineage. Nevertheless, it is clear that the *cpe* cluster in *M. aeruginosa* has originated from a single ancestor, and was introduced into *M. aeruginosa* early during evolution.

**Fig. 5.**
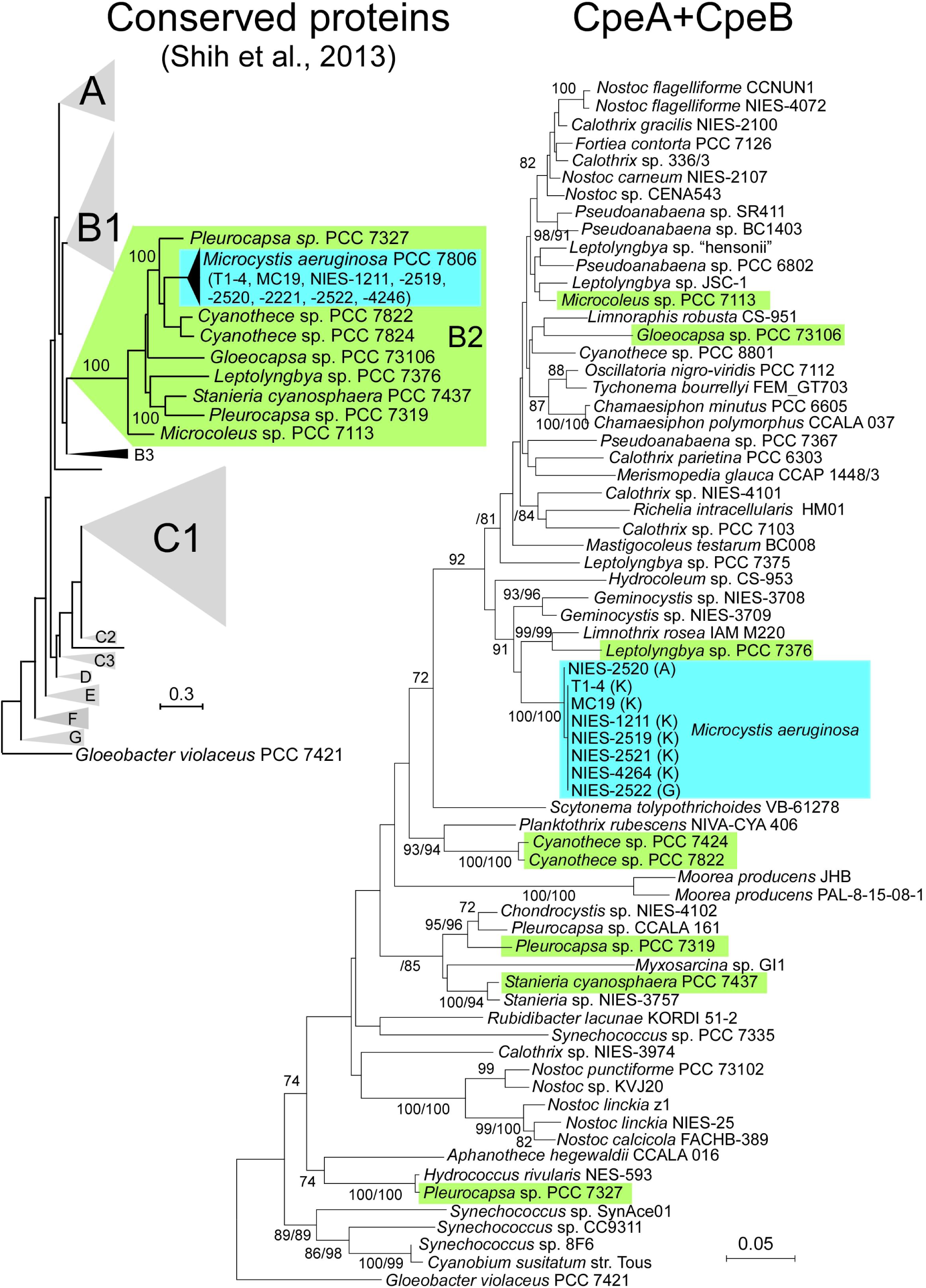
Molecular phylogenetic trees of *Cyanobacteria*. **a,** The phylogenetic tree modified from the conserved protein-based tree by Shih et al., (2013). RaxML bootstrap values (>90) are indicated at the respected nodes. All PE-containing *M. aeruginosa* strains were not included in the analysis. However, whole-genome blast analyses indicated that all these strains are most closely related to PCC 7806 (Frangeul et al., 2008) in the tree (indicated in parentheses). **b**, A NJ phylogenetic tree of concatenated amino acid sequences of CpeA and CpeB (336 amino acids in total). Bootstrap values (NJ >70/ RAxML >80) on the basis of 1 000 replicates are indicated at the respective nodes. Numbers in parentheses after the strain name of *M. aeruginosa* indicate group assignment. The sequence of *Gloeobacter violaceus* PCC 7421 is used as an outgroup as in **a**. The color-coding in **b** is according to **a**. Scale bar, 0.05 substitutions per sites.

**Fig. 6.**
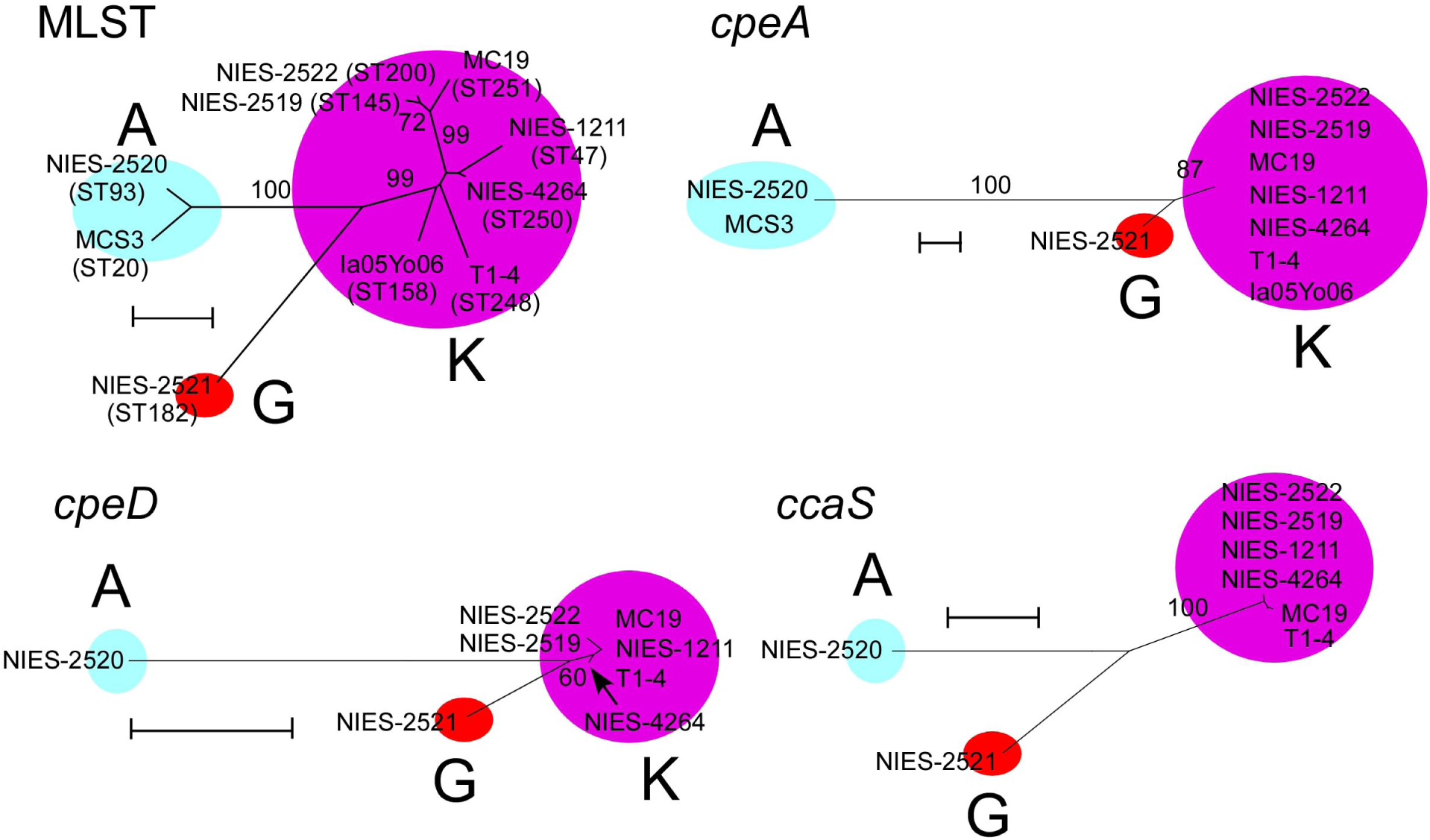
Molecular phylogenetic trees of *M. aeruginosa*. NJ trees on the basis of nucleotide sequence of concatenation of the seven MLST loci (**a**) (using only PE-containing strains), *cpeA* (**b**) (495 bps), *cpeD* (**c**) (768 bps), *ccaS* (**d**) (2 694 bps). Scale bar, 0.005 substitutions per sites.

**Fig. 7.**
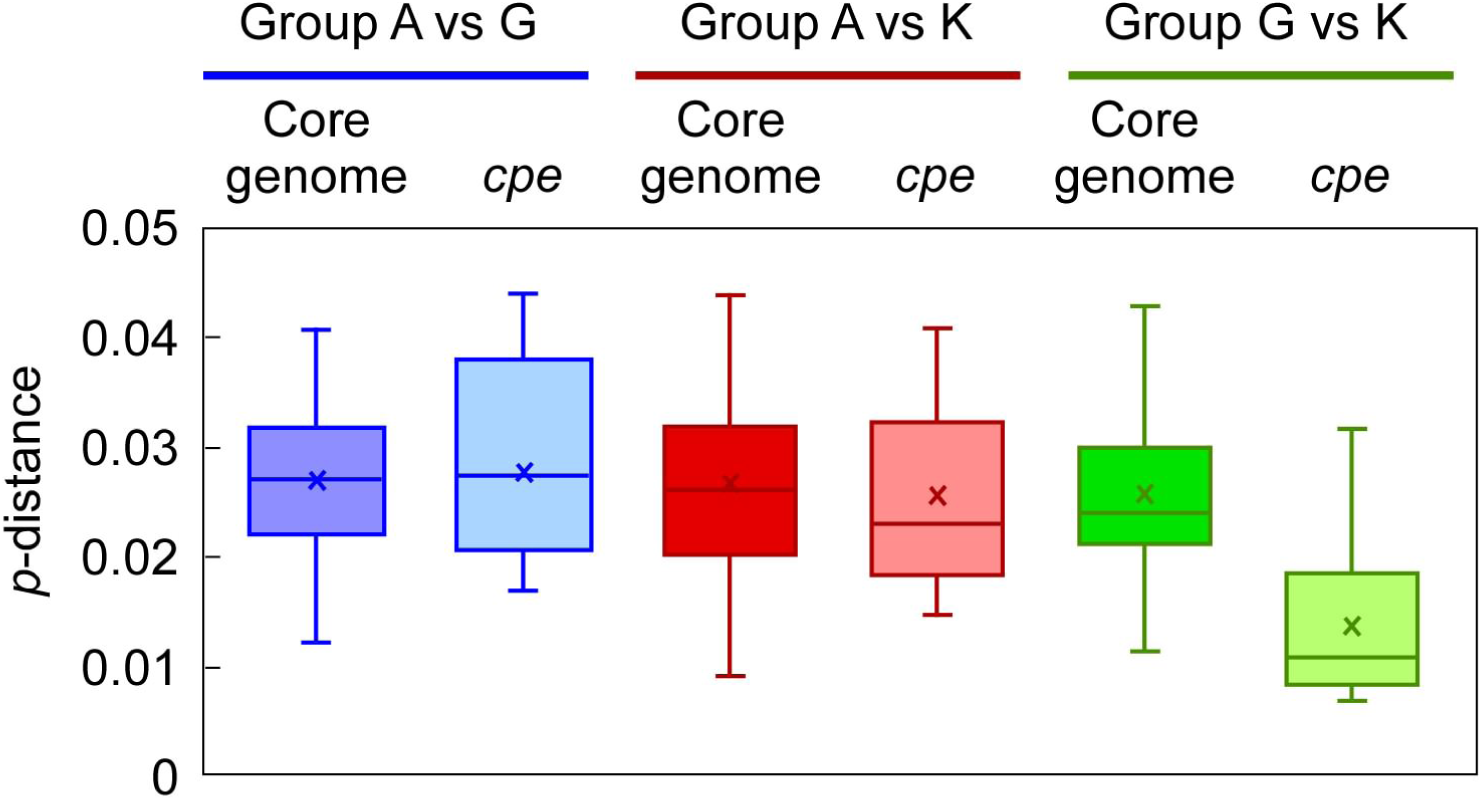
A Box-whisker plot of nucleotide divergence. Y-axis indicates percent nucleotide divergence (*p*-distance, calculated using MEGA) between the two groups. For distance calculation, NIES-1211 is used as the representative of group K. Core genome genes used for the comparison included 71 housekeeping genes including the seven MLST genes, and *mutS2* which is adjacent to the *cpe* cluster (Supplementary Table S4). PE genes included 12 PE-containing strain-specific genes (indicated in the red box in Figure 4). *cpeR* was not included because of low nucleotide divergence, probably owing to its short length (333 bps).

**Fig. 8.**
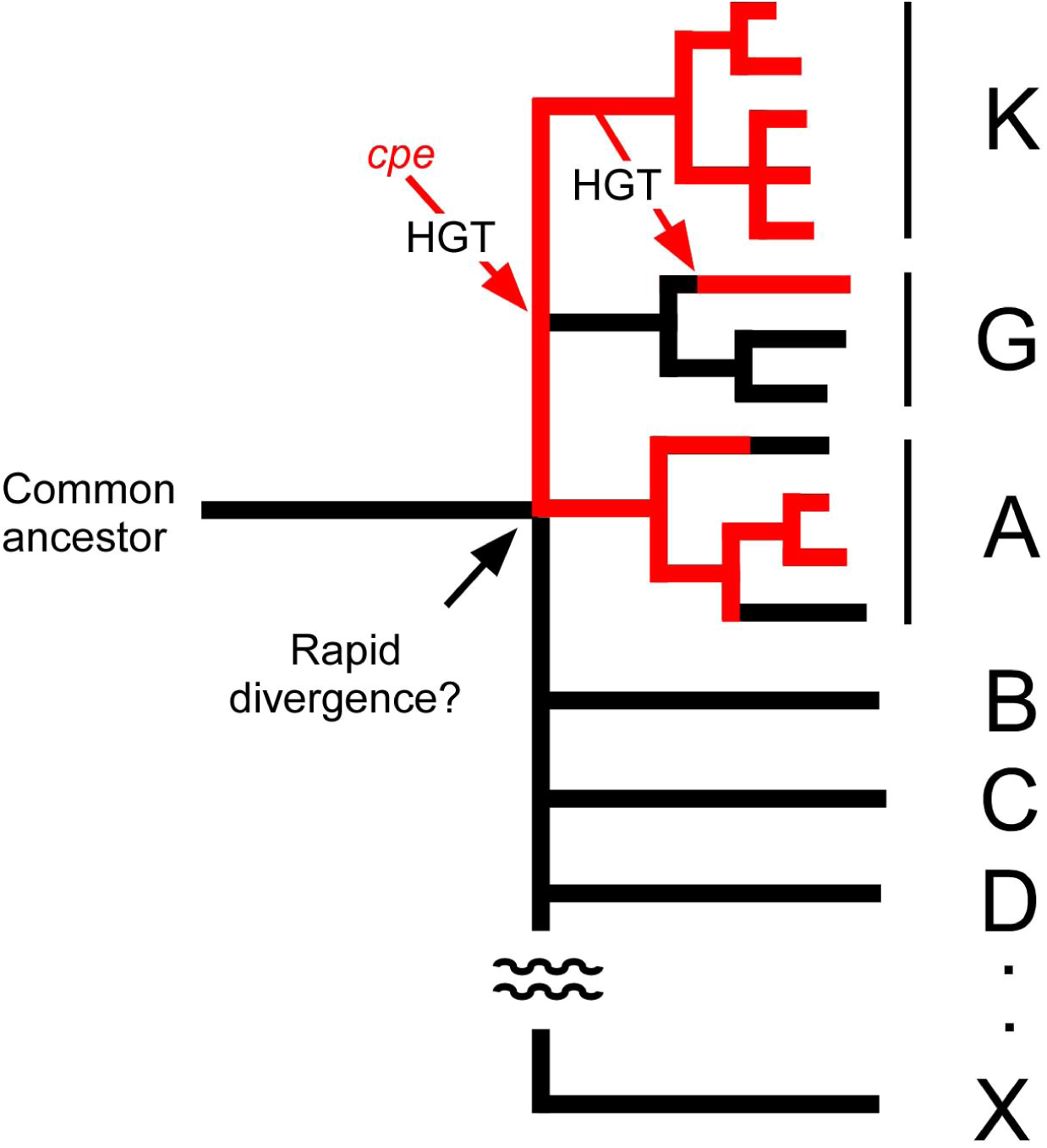
A schematic representation of the possible evolutionary history of PE genes in *M. aeruginosa*. Red lines indicate lineages containing PE genes. Alphabets indicate major intraspecies groups defined in Figure 3. The branching orders of these groups are not clear due to low statistical resolution (Figure 3), which might reflect a rapid divergence of these groups.

### Ecological and evolutionary implication of PE pigmentation in *M. aeruginosa*

The current data suggest that PE-containing strains are divided into three phylogenetic groups. In particular, group K isolates are the most abundant PE-positive genotypes and most probably represent a stable and long-lasting ecotype (Cohan, 2002). This suggests that PE contained in the phycobilisome presents some ecological advantage over PE-negative strains of *M. aeruginosa* in ecological niches. A classic consideration points to the advantage of PE over non-PE pigmentation; deep underwater environments, where light of longer wavelength is limited, are niches where an effective application of green light is necessary (Kirk, 1994). However, *M. aeruginosa* does not appear to favor deepwater for green light and is known to perform a vertical movement from the bottom for nutrition to the water surface for light energy. This is performed by using the balance between expression of the gas vesicle protein (Gvp) and photosynthetic carbohydrate accumulation inside the cell (Reynolds et al., 1987). Indeed, the five PE-containing strains (NIES-2519, NIES-2520, NIES-2521, NIES-2522, and NIES4264) were isolated from surface waters and BLAST analyses showed that all PE-containing *M. aeruginosa* harbor *gvp* that encode Gvp proteins (Mlouca et al., 2004). These observations exclude the possible preference of PE-containing strains for deepwater. The PE pigmentation *in M. aeruginosa* can reasonably be explained by a more recent theory, wherein the portioning of the light spectrum enables a coexistence of phytoplanktons with preference for different wavelengths and thereby driving adaptive pigment divergence in phytoplanktons (Stomp et al., 2004). This theory has been supported by studies on natural water study and has suggested that intermediate turbidity favors the coexistence of different pigment types even in shallow water (Stomp et al., 2007). This is indeed consistent with the frequent coexistence of both PE-positive and negative genotypes in the same bloom of *M. aeruginosa.* For example, NIES-2521 and other PE-negative group G strains have been co-isolated from the same bloom sample in eutrophic shallow water, Lake Kasumigaura where the maximum depth is 7 m (Tanabe et al., 2009). Extensive field survey of the spatial distribution of PE-containing *M. aeruginosa* strains together with laboratory co-culture experiments would further clarify the coexistence of PE-positive and negative strains. In the context of evolutionary history, the observed coexistence of PE-positive and -negative strains of *M. aeruginosa* is a likely result of the adaptive radiation via interspecific HGT of PE genes. Similar evolutionary patterns have been reported for other cyanobacteria such as *Synechococcus* spp. (Haverkamp et al., 2009; Six et al., 2007).

## Conclusions

Whole genome analyses identified the *cpe* cluster in PE-containing strains of *M. aeruginosa* among different lineages. All PE-containing species adjust PE but not PC contents under different green/red light regime, representing a type II CA. The *cpe* cluster of *M. aeruginosa* has a single origin, and has been imported from other cyanobacterial species via HGT which possibly occurred prior to the divergence of major intraspecies groups, A and K. Each PE-containing phylogenetic cluster probably represents a stable ecotype, which has likely adapted to an ecological niche where green light is accessible. The future identification and characterization of PE-containing strains with diverse genomic backgrounds would further clarify the diversity and evolution of PE pigmentation and its regulation in *M. aeruginosa*.

## Supporting information

## Acknowledgements

We thank Yuu Hirose for his invaluable advice and discussion regarding the data interpretation, Iwane Suzuki and Makoto M. Watanabe for the use of their laboratory facilities, and Nobuyoshi Nakajima and Shigekatsu Suzuki for technical assistance of the sequencing. This work was financially supported by JSPS KAKENHI Grant Number 15K07523 and 16H02943.

## Author contributions

YT designed the research, isolated strains, performed whole genome analyses, phylogenetic analyses, and pigment analyses, HY performed whole genome shotgun sequencing, and YT and HY wrote the paper. All authors read and approved the final manuscript.

## References

Bankevich, A., Nurk, S., Antipov, D., Gurevich, A. A., Dvorkin, M., Kulikov, A. S., et al. (2012). SPAdes: a new genome assembly algorithm and its applications to single-cell sequencing. J. Comput. Biol. 19, 455–477. doi:10.1089/cmb.2012.0021

Beasley, V. R., Cook, W. O., Dahlem, A. M., Hooser, S. B., Lovell, R. A., and Valentine, W. M. (1989). Algae intoxication in livestock and waterfowl. Food Anim. Pract. 5, 345–361. doi:10.1016/S0749-0720(15)30980-4

Cohan, F. M. (2002). What are bacterial species? Annu. Rev. Microbiol. 56, 457–487. doi:10.1146/annurev.micro.56.012302.160634

Edgar, R. C. (2004). MUSCLE: multiple sequence alignment with high accuracy and high throughput. Nucleic Acids Res. 32, 1792–1797. doi:10.1093/nar/gkh340

Frangeul, L., Quillardet, P., Castets, A. M., Humbert, J. F., Matthijs, H. C., Cortez, D., et al. (2008). Highly plastic genome of *Microcystis aeruginosa* PCC 7806, a ubiquitous toxic freshwater cyanobacterium. BMC Genomics 9, 1. doi:10.1186/1471-2164-9-274

Guljamow, A., Jenke-Kodama, H., Saumweber, H., Quillardet, P., Frangeul, L., Castets, A. M., et al. (2007). Horizontal gene transfer of two cytoskeletal elements from a eukaryote to a cyanobacterium. Curr. Biol. 17, R757–R759. doi:10.1016/j.cub.2007.06.063

Harke, M. J., Steffen, M. M., Gobler, C. J., Otten, T. G., Wilhelm, S. W., Wood, S. A., et al. (2016). A review of the global ecology, genomics, and biogeography of the toxic cyanobacterium, *Microcystis* spp. Harmful Algae 54, 4–20. doi:10.1016/j.hal.2015.12.007

Haverkamp, T. H., Schouten, D., Doeleman, M., Wollenzien, U., Huisman, J., and Stal, L. J. (2009). Colorful microdiversity of *Synechococcus* strains (picocyanobacteria) isolated from the Baltic Sea. ISME J. 3, 397–408. doi:10.1038/ismej.2008.118

Hirose, Y., Misawa, N., Yonekawa, C., Nagao, N., Watanabe, M., Ikeuchi, M.. et al. (2017). Characterization of the genuine type 2 chromatic acclimation in the two *Geminocystis* cyanobacteria. DNA Res. 24, 387–396. doi:10.1093/dnares/dsx011

Hirose, Y., Narikawa, R., Katayama, M., and Ikeuchi, M. (2010). Cyanobacteriochrome CcaS regulates phycoerythrin accumulation in *Nostoc punctiforme*, a group II chromatic adapter. Proc. Natl. Acad. Sci. USA 107, 8854–8859. doi:10.1073/pnas.1000177107

Humbert, J. F., Barbe, V., Latifi, A., Gugger, M., Calteau, A., Coursin, T., et al. (2013). A tribute to disorder in the genome of the bloom-forming freshwater cyanobacterium *Microcystis aeruginosa*. PLoS ONE 8, e70747. doi:10.1371/journal.pone.0070747

Ikeuchi, M., and Ishizuka, T. (2008). Cyanobacteriochromes: a new superfamily of tetrapyrrole-binding photoreceptors in cyanobacteria. Photochem. Photobiol. Sci. 7, 1159–1167. doi:10.1039/B802660M

Jeong, H., Chun, S. J., Srivastava, A., Cui, Y., Ko, S. R., Oh, H. M., et al. (2018). Genome sequences of two cyanobacterial strains, toxic green *Microcystis aeruginosa* KW (KCTC 18162P) and nontoxic brown *Microcystis* sp. Strain MC19, under xenic culture conditions. Genome Announc. 6, e00378–18. doi:10.1128/genomeA.00378-18

Jochimsen, E. M., Carmichael, W. W., An, J., Cardo, D. M., Cookson, S. T., Holmes, C. E., et al. (1998). Liver failure and death after exposure to microcystins at a hemodialysis center in Brazil. New Engl. J. Med. 338, 873–878. doi:10.1056/NEJM199803263381304

Kasai, F., Kawachi, M., Erata, M., and Watanabe, M. M. (2004). NIES-Collection, List of strains, microalgae and protozoa, 7th edn. Tsukuba, Japan: National Institute for Environmental Studies.

Kirk, J. T. O. (1994). Light and photosynthesis in aquatic ecosystems. Cambridge: Cambridge University Press.

Miller, M. A., Kudela, R. M., Mekebri, A., Crane, D., Oates, S. C., Tinker, M. T., et al. (2010a). Evidence for a novel marine harmful algal bloom: cyanotoxin (microcystin) transfer from land to sea otters. PLoS ONE5, e12576. doi:10.1371/journal.pone.0012576

Miller, M. A., Pfeiffer, W., and Schwartz, T. (2010b). “Creating the CIPRES Science Gateway for inference of large phylogenetic trees” in Proceedings of the Gateway Computing Environments Workshop (GCE), 14 Nov. 2010, New Orleans, 1–8.

Mlouka, A., Comte, K., Castets, A. M., Bouchier, C., and de Marsac, N. T. (2004). The gas vesicle gene cluster from *Microcystis aeruginosa* and DNA rearrangements that lead to loss of cell buoyancy. J. Bacteriol. 186, 2355–2365. doi:10.1128/JB.186.8.2355-2365.2004

Montgomery, B. L. (2017). Seeing new light: recent insights into the occurrence and regulation of chromatic acclimation in cyanobacteria. Curr. Opin. Plant Biol. 37, 18–23. doi:10.1016/j.pbi.2017.03.009

Otsuka, S., Suda, S., Li, R., Watanabe, M., Oyaizu, H., Hiroki, M., et al. (1998a). Phycoerythrin‐containing *Microcystis* isolated from PR China and Thailand. Phycol. Res. 46, 45–50. doi:10.1046/j.1440-1835.1998.00124.x

Otsuka, S., Suda, S., Li, R., Watanabe, M., Oyaizu, H., Matsumoto, S., et al. (1998b). 16S rDNA sequences and phylogenetic analyses of Microcystis strains with and without phycoerythrin. FEMS Microbiol. Lett. 164, 119–124. doi:10.1111/j.1574-6968.1998.tb13076.x

Otsuka, S., Suda, S., Li, R., Watanabe, M., Oyaizu, H., Matsumoto, S., et al. (1999). Phylogenetic relationships between toxic and non-toxic strains of the genus *Microcystis* based on 16S to 23S internal transcribed spacer sequence. FEMS Microbiol. Lett. 172, 15–21. doi:10.1111/j.1574-6968.1999.tb13443.x

Reynolds, C. S., Oliver, R. L., and Walsby, A. E. (1987). Cyanobacterial dominance: the role of buoyancy regulation in dynamic lake environments. New Zeal. J. Mar. Fresh. 21, 379–390. doi:10.1080/00288330.1987.9516234

Schatz, D., Keren, Y., Vardi, A., Sukenik, A., Carmeli, S., Börner, T., et al. (2007). Towards clarification of the biological role of microcystins, a family of cyanobacterial toxins. Environ. Microbiol. 9, 965–970. doi:10.1111/j.1462-2920.2006.01218.x

Seemann, T. (2014). Prokka: rapid prokaryotic genome annotation. Bioinformatics 30, 2068–2069. doi:10.1093/bioinformatics/btu153

Shih, P. M., Wu, D., Latifi, A., Axen, S. D., Fewer, D. P., Talla, E., et al. (2013). Improving the coverage of the cyanobacterial phylum using diversity-driven genome sequencing. Proc. Natl. Acad. Sci. USA 110, 1053–1058. doi:10.1073/pnas.1217107110

Six, C., Thomas, J. C., Garczarek, L., Ostrowski, M., Dufresne, A., Blot, N., et al. (2007). Diversity and evolution of phycobilisomes in marine *Synechococcus* spp.: a comparative genomics study. Genome Biol. 8, R259. doi:10.1186/gb-2007-8-12-r259

Stamatakis, A. (2006). RAxML-VI-HPC: maximum likelihood-based phylogenetic analyses with thousands of taxa and mixed models. Bioinformatics 22, 2688–2690. doi:10.1093/bioinformatics/btl446

Stomp, M., Huisman, J., De Jongh, F., Veraart, A. J., Gerla, D., Rijkeboer, M., et al. (2004). Adaptive divergence in pigment composition promotes phytoplankton biodiversity. Nature 432, 104–107. doi:10.1038/nature03044

Stomp, M., Huisman, J., Vörös, L., Pick, F. R., Laamanen, M., Haverkamp, T., et al. (2007). Colourful coexistence of red and green picocyanobacteria in lakes and seas. Ecol. Lett. 10, 290–298. doi:10.1111/j.1461-0248.2007.01026.x

Tamura, K., Stecher, G., Peterson, D., Filipski, A., and Kumar, S. (2013) MEGA6: molecular evolutionary genetics analysis version 6.0. Mol. Biol. Evol. 30, 2725–2729. doi:10.1093/molbev/mst197

Tanabe, Y., Hodoki, Y., Sano, T., Tada, K., and Watanabe, M. M. (2018). Adaptation of the freshwater bloom-forming cyanobacterium *Microcystis aeruginosa* to brackish water is driven by recent horizontal transfer of sucrose genes. Front. Microbiol. 9, 1150. doi:10.3389/fmicb.2018.01150

Tanabe, Y., Kasai, F., and Watanabe, M. M. (2007). Multilocus sequence typing (MLST) reveals high genetic diversity and clonal population structure of the toxic cyanobacterium *Microcystis aeruginosa*. Microbiology 153, 3695–3703. doi:10.1099/mic.0.2007/010645-0

Tanabe, Y., Kasai, F., and Watanabe, M. M. (2009). Fine-scale spatial and temporal genetic differentiation of water bloom-forming cyanobacterium *Microcystis aeruginosa*: revealed by multilocus sequence typing. Environ. Microb. Rep. 1, 575–582. doi:10.1111/j.1758-2229.2009.00088.x

Tanabe, Y., and Watanabe, M. M. (2011). Local expansion of a panmictic lineage of water bloom-forming cyanobacterium *Microcystis aeruginosa*. PLoS ONE 6, e17085. doi:10.1371/journal.pone.0017085

Tandeau De Marsac, N. (1977). Occurrence and nature of chromatic adaptation in cyanobacteria. J. Bacteriol. 130, 82–91.

Tillett, D., Dittmann, E., Erhard, M., von Döhren, H., Börner, T., and Neilan, B. A. (2000). Structural organization of microcystin biosynthesis in *Microcystis aeruginosa* PCC7806: an integrated peptide–polyketide synthetase system. Chem. Biol. 7, 753–764. doi:10.1016/S1074-5521(00)00021-1

Tooming-Klunderud, A., Sogge, H., Rounge, T. B., Nederbragt, A. J., Lagesen, K., Glöckner, G. et al. (2013). From green to red: horizontal gene transfer of phycoerythrin gene cluster between *Planktothrix* strains. Appl. Environ. Microbiol. 79, 6803–6812. doi:10.1128/AEM.01455-13

Watanabe, M., and Ikeuchi, M. (2013). Phycobilisome: architecture of a light-harvesting supercomplex. Photosynth. Res. 116, 265–276. doi:10.1007/s11120-013-9905-3

