## Supplementary material for "Evolutionary history of phycoerythrin pigmentation in the water bloom-forming cyanobacterium *Microcystis aeruginosa*"

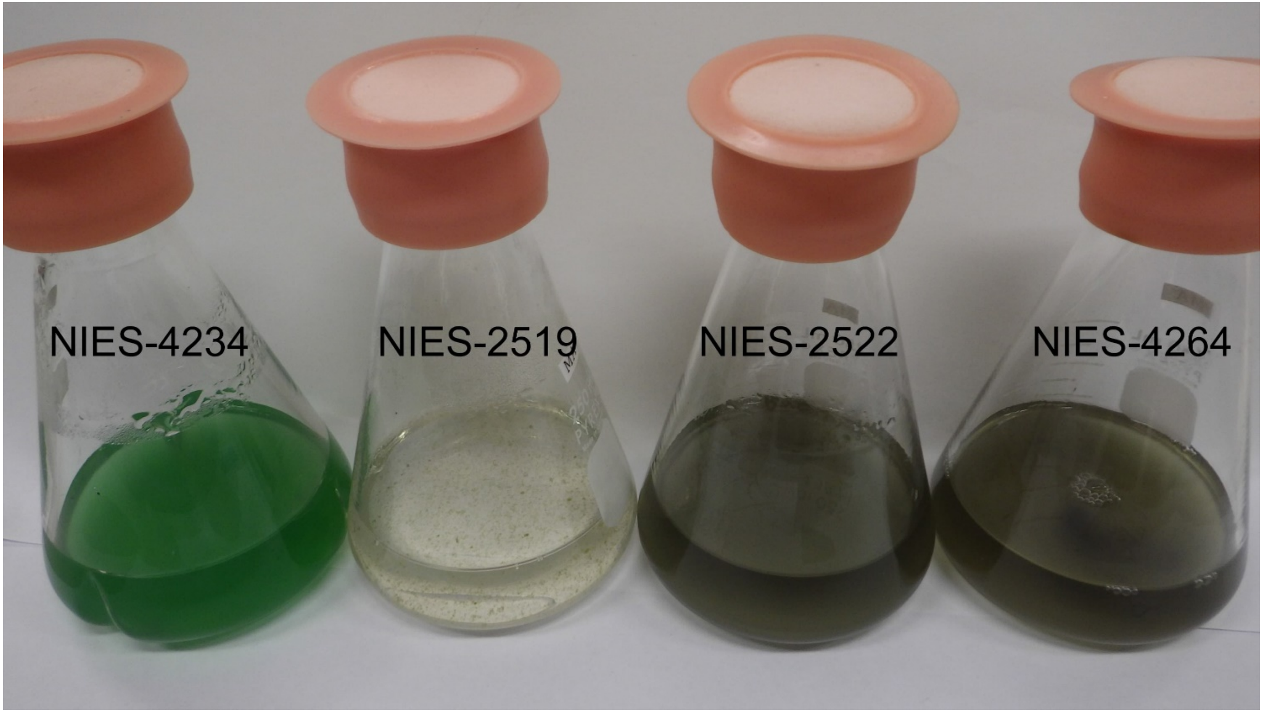


**Supplementary Figure S1.** Cultures of *M. aeruginosa* strain grown under white light. NIES-2519, NIES-2522, and NIES-4264 are PE-containing strains, whereas NIES-4234 is the PE-deficient strain (control).

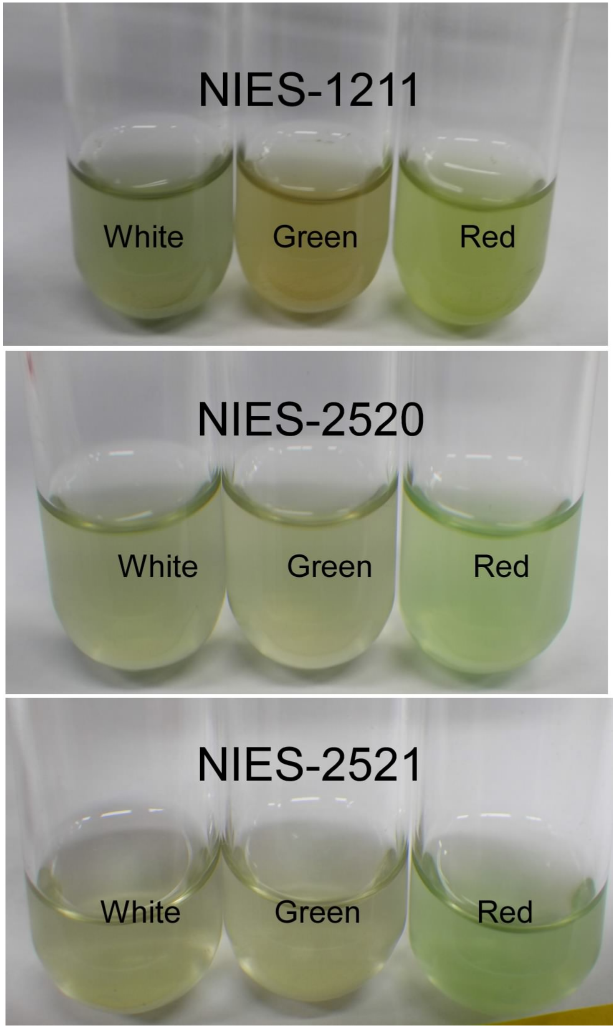


**Supplementary Figure S2.** Cultures of PE-containing *M. aeruginosa* strains grown under white, green, and red light. Note that all strains turned green in color under red light, indicative of chromatic adaptation (CA).

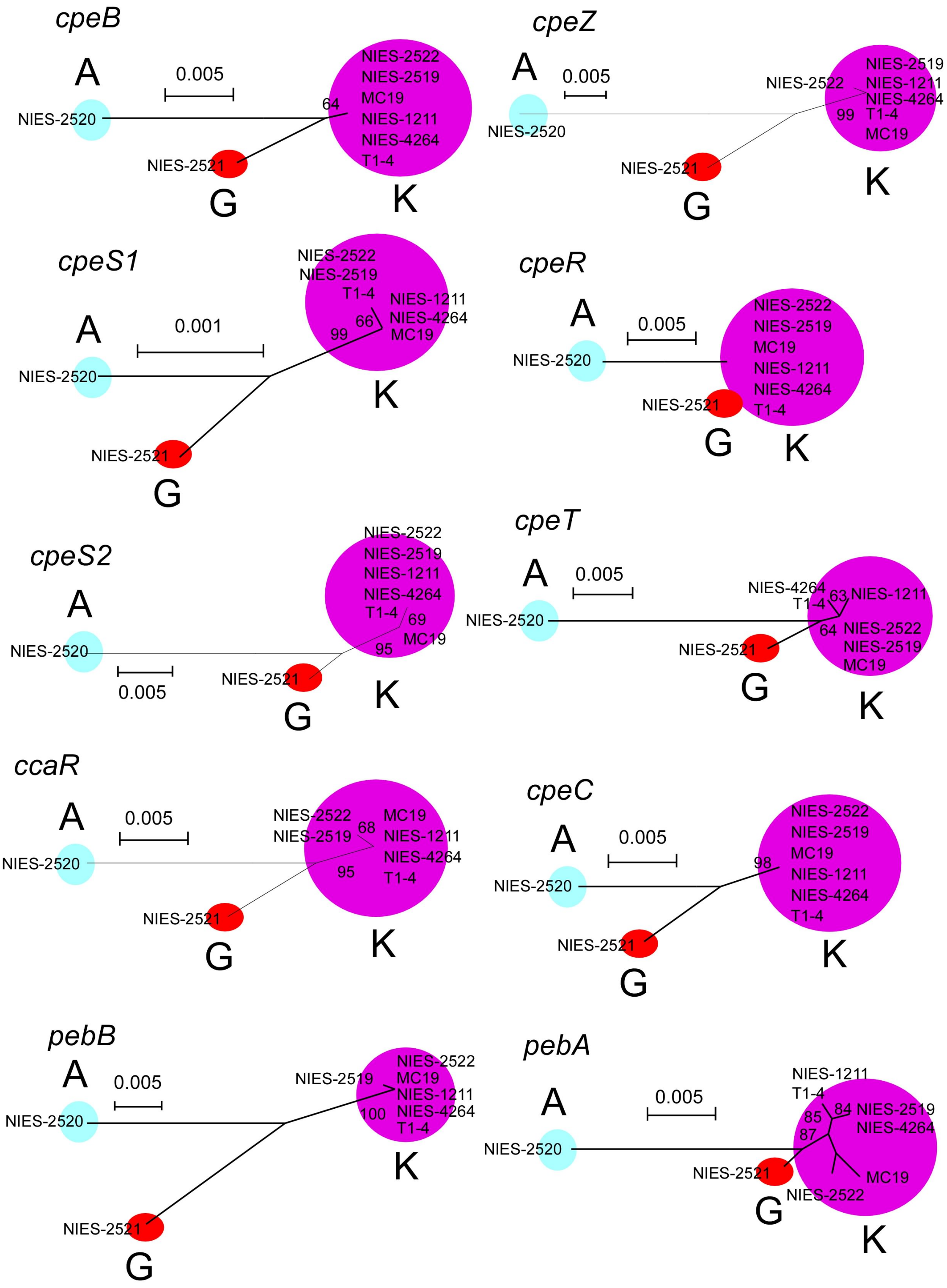


**Supplementary Figure S3.** NJ tress of PE genes. NJ bootstrap values were indicated at the respective nodes. Note that all trees showed the same topology regarding the relationships among groups A, G, and K. Scale bars, substitutions per site.

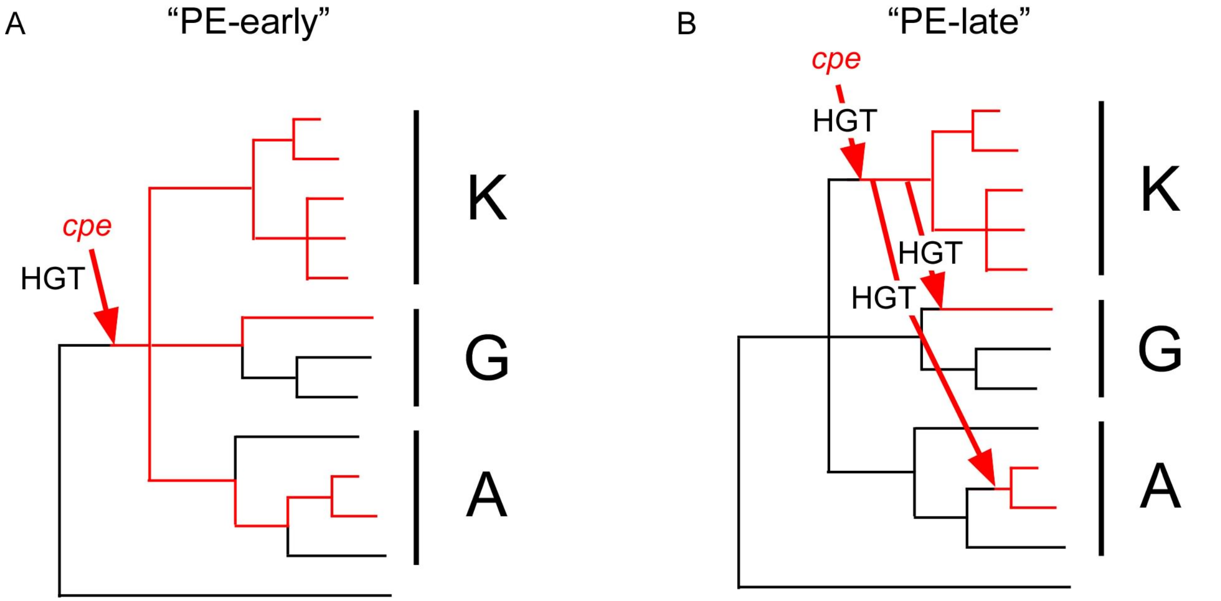


**Supplementary Figure S4.** Schematic representation of the two possible scenarios of PE gene evolution in *M. aeruginosa*. **(A)**, The "PE-early" hypothesis suggests that one *cpe* gene cluster is imported from other species and is repeatedly lost thereafter with no HGT. **(B)**, The "PE-late" hypothesis suggests that one *cpe* gene cluster is imported, followed by multiple HGTs into different lineages of *M. aeruginosa*.
